# AlphaFill: enriching the AlphaFold models with ligands and co-factors

**DOI:** 10.1101/2021.11.26.470110

**Authors:** Maarten L. Hekkelman, Ida de Vries, Robbie P. Joosten, Anastassis Perrakis

**Author notes:** These authors have contribute equally to the work reported (Maarten L. Hekkelman, Ida de Vries). These authors have jointly supervised the project (Robbie P. Joosten, Anastassis Perrakis).

## Abstract

Artificial intelligence (AI) methods for constructing structural models of proteins on the basis of their sequence are having a transformative effect in biomolecular sciences. The AlphaFold protein structure database makes available hundreds of thousands of protein structures. However, all these structures lack cofactors essential for their structural integrity and molecular function (e.g. hemoglobin lacks a bound heme), key ions essential for structural integrity (e.g. zinc-finger motifs) or catalysis (e.g. Ca^2+^ or Zn^2+^ in metalloproteases), and ligands that are important for biological function (e.g. kinase structures lack ADP or ATP). Here, we present AlphaFill, an algorithm based on sequence and structure similarity, to “transplant” such “missing” small molecules and ions from experimentally determined structures to predicted protein models. These publicly available structural annotations are mapped to predicted protein models, to help scientists interpret biological function and design experiments.

## Introduction

The protein folding problem, in practical terms predicting a protein 3D structure based on only its amino acid sequence, has been a challenge in structural biology for decades. Recently, artificial intelligence (AI) approaches have made protein structure prediction possible, as implemented in the AlphaFold^1^ and the RoseTTAfold^2^ methods. Both approaches create impressively accurate structure models at the domain level; flexible parts of the protein (such as loops) are predicted with lower accuracy and confidence. The structure predictions for the proteomes of ten different organisms are already publicly available in the AlphaFold protein structure database^3^, which currently contains 365,198 predicted protein structures, and more will follow. These predicted models can provide new biological insights with regard to protein function, particularly for structures that have not been determined experimentally.

An interesting feature of the AI prediction algorithms is that in reality they have not learned to solve the protein folding problem from the perspective of understanding “first principles” of how protein folding occurs in nature: they have merely but impressively learned to circumvent it based on known experimental structures. Consequently, many of the well-predicted models cannot occur in nature: myoglobin or hemoglobin need a heme to fold, zinc-finger structures cannot be stable without a zinc ion, and many proteins can only exist as homo- or hetero-multimers. However, the structural models in the AlphaFold database disregard all chemical entities other than natural amino-acid residues that are needed for the folding or the function of many proteins and do not contain multimers or complexes with other proteins^4^. The latter issue, is now being circumvented by the development of AlphaFoldMultimer^5^ and RoseTTAFold^6^, that are able to predict biological assemblies that are provided by the user at the sequence and stoichiometry level.

Here, we enrich the models in the AlphaFold database by “transplanting” several common small molecules and ions that have been observed in complex with highly similar homologous proteins in experimentally determined models from the PDB-REDO databank^7^. We present the AlphaFill algorithm and apply it to all AlphaFold models, to make a new resource: the AlphaFill databank. We describe this new resource and discuss examples illustrating how it can help scientists to better understand and use predicted structure models to answer relevant scientific questions.

## Results

### A procedure for “transplanting” small molecules and ions to AlphaFold models

The *AlphaFill* procedure, for filling-up missing information to AlphaFold models, goes through the following steps.

- The sequence of the AlphaFold model is BLASTed^8^ against the sequence file of the LAHMA webserver^9^ which contains all sequences present in the PDB-REDO databank. The hits are sorted by E-value and a maximum of 250 hits, as is the default for BLAST, is returned.
- All hits are retrieved from the PDB-REDO databank and checked for compounds of interest for AlphaFill (*vide infra*).
- The remaining hits are filtered to ensure only sufficiently close homologs are used: consistent with our other software that employs homology^7,9,10^, the sequence identity cutoff is currently 70% over an aligned sequence of at least 85 residues. The latter cut-off was assigned empirically to filter for domain-sized units.
- The selection of hits is then structurally aligned^11^, based on the Cα-atoms of the residues matched in the BLAST^8^ alignment. The root-mean-square deviation (RMSD) of this global alignment is stored in the AlphaFill metadata.
- Starting from the hit with the best E-value, each compound of interest (see below) in the hit list is scanned for its local surroundings. All backbone atoms within 6Å are then used for a local structural alignment to the AlphaFold model. The RMSD of this local alignment is stored in the AlphaFill metadata.
- Compounds are then integrated into the AlphaFold model, unless the same compound is already placed within 3.5Å of the centroid of the compound to be fitted. If compounds have multiple alternate conformations, all conformations are transferred. Descriptions of covalent bonds or metal binding captured in so-called *struct_conn* records are also integrated in the AlphaFill model.
- The final AlphaFill model with all transplanted compounds is then stored as mmCIF coordinate file together with a JSON-formatted metadata file describing the provenance of each transplanted compound.

The running time per model depends strongly on the number of BLAST hits and compounds to be transferred. Based on calculations with all AlphaFold models, the mean running time is 2 minutes per model on a single Intel Xeon E5-2697 CPU thread.

### A collection of biologically relevant co-factors, small molecule ligands and ions

To create AlphaFill models that contain the relevant compounds to represent a biological state suited for further study, a collection of cofactors, ligands and metal ions was created (see Methods for details on how this collection was constructed). Briefly, a list of the most common compounds in the PDB was constructed and supplemented by the cofactors and their analogues listed in the CoFactor database^12^. Where possible, cofactor analogues were mapped to the representative co-factor by atom renaming (and atom deletion), e.g. ACP (Adenosine-5’-[beta,gamma-methylene]triphosphate; methylene substituted ATP) is mapped to ATP, as ATP is the compound involved in biological processes. Cofactor adducts like CNC (Vitamin B12 in complex with cyanide) are trimmed down to their parent cofactor (e.g. Vitamin B12 in the CNC case) by atom deletion. Common crystallisation agents (e.g. poly-ethyleneglycol and chloride), compounds used for phasing (e.g. cadmium ions) and carbohydrates were purposely excluded. The current collection of compounds to be transplanted consists of 400 entries. It is stored as a cif-formatted file separate from the *AlphaFill* program to allow easy extension in future incarnations of the AlphaFill databank. This cif-file is available through the AlphaFill databank.

### The AlphaFill databank

Applying *AlphaFill* to the current version of the AlphaFold database (November 2021, 365,198 models) resulted in 21,899 models that had at least one transplanted compound. A total of 108,997 compounds were transplanted in these models. A selection of frequently transplanted compounds is listed in Table 1, including their “transplantation” frequency. A full table of all transplanted compounds is available as Supplemental Table S1. All AlphaFill models are available from https://alphafill.eu through a web-based user interface. In addition, the whole databank can be downloaded by means of rsync through rsync.alphafill.eu. This also includes all relevant metadata: the JSON description of all the transplants for each AlphaFill model, a JSON schema with a complete description of these files, and the current cif-file that describes the compounds that are considered for transfer.

**Table 1:** Examples of frequently transplanted compounds in the AlphaFill databank.

| Compound | Description | # entries | # transplants |
| --- | --- | --- | --- |
| **Nucleotides** |  |  |  |
| ADP | Adenosine diphosphate | 2198 | 2928 |
| GDP | Guanosine diphosphate | 1655 | 2468 |
| ATP | Adenosine triphosphate | 1221 | 1517 |
| AMP | Adenosine monophosphate | 723 | 926 |
| GTP | Guanosine triphosphate | 686 | 921 |
| UDP | Uridine diphosphate | 243 | 267 |
| **Cofactors** |  |  |  |
| NAD | Nicotinamide adenine dinucleotide | 685 | 978 |
| HEM | Heme | 498 | 658 |
| FAD | Flavin adenine dinucleotide | 470 | 628 |
| NAP | NAD phosphate/NADP | 455 | 618 |
| PLP | Pyridoxal phosphate | 369 | 602 |
| SAH | S-Adenosyl-L-homocysteine | 355 | 383 |
| GSH | Glutathione | 324 | 450 |
| NDP | NADPH | 317 | 445 |
| SAM | S-adenosylmethionine | 298 | 407 |
| COA | Coenzyme A | 254 | 368 |
| NAI | NADH | 204 | 298 |
| FMN | Flavin mononucleotide | 197 | 266 |
| **Miscellaneous** |  |  |  |
| CLA | Chlorophyll A | 163 | 4058 |
| CLR | Cholesterol | 157 | 380 |
| **(Metal) ions** |  |  |  |
| MG | Magnesium ion | 8069 | 19754 |
| NA | Sodium ion | 5650 | 15262 |
| ZN | Zinc ion | 5261 | 13051 |
| CA | Calcium ion | 5078 | 17312 |
| K | Potassium ion | 2267 | 3910 |

### A web-based user interface for the AlphaFill databank

All AlphaFill entries are available for visual inspection through the AlphaFill website. In the front page models can be retrieved using the AlphaFold identifier. Individual entries can also be pointed to directly using the AlphaFold identifier, e.g. https://alphafill.eu/?id=P02144 for human myoglobin. The web site also makes available the compound prevalence (on the *Compounds* page), as well as numbers regarding transplanted compounds for each “filled” AlphaFold model (on the *Structures* page).

On each entry page (Figure 1) the model is displayed using Mol*^13^, allowing users full flexibility for inspection. By default, all transplanted compounds are shown, listed by compound and sorted by increasing RMSD (global at the hit level and local at the individual compound level). These compounds are also listed in a table together with the parent PDB-REDO entry, the global Cα alignment RMSD, the name of the compound (plus the original name if it was mapped), and the local alignment RMSD. Clicking a row in the table changes the focus of the viewer to that compound. Compounds can also be toggled on and off to reduce clutter in the view. Transplants based on models with a Cα alignment RMSD > 5.0Å or a local alignment RMSD > 1.0Å are considered low confidence and are marked in the table with a red background. The model with all the ligands and the metadata can be downloaded for further inspection.

**Figure 1:**
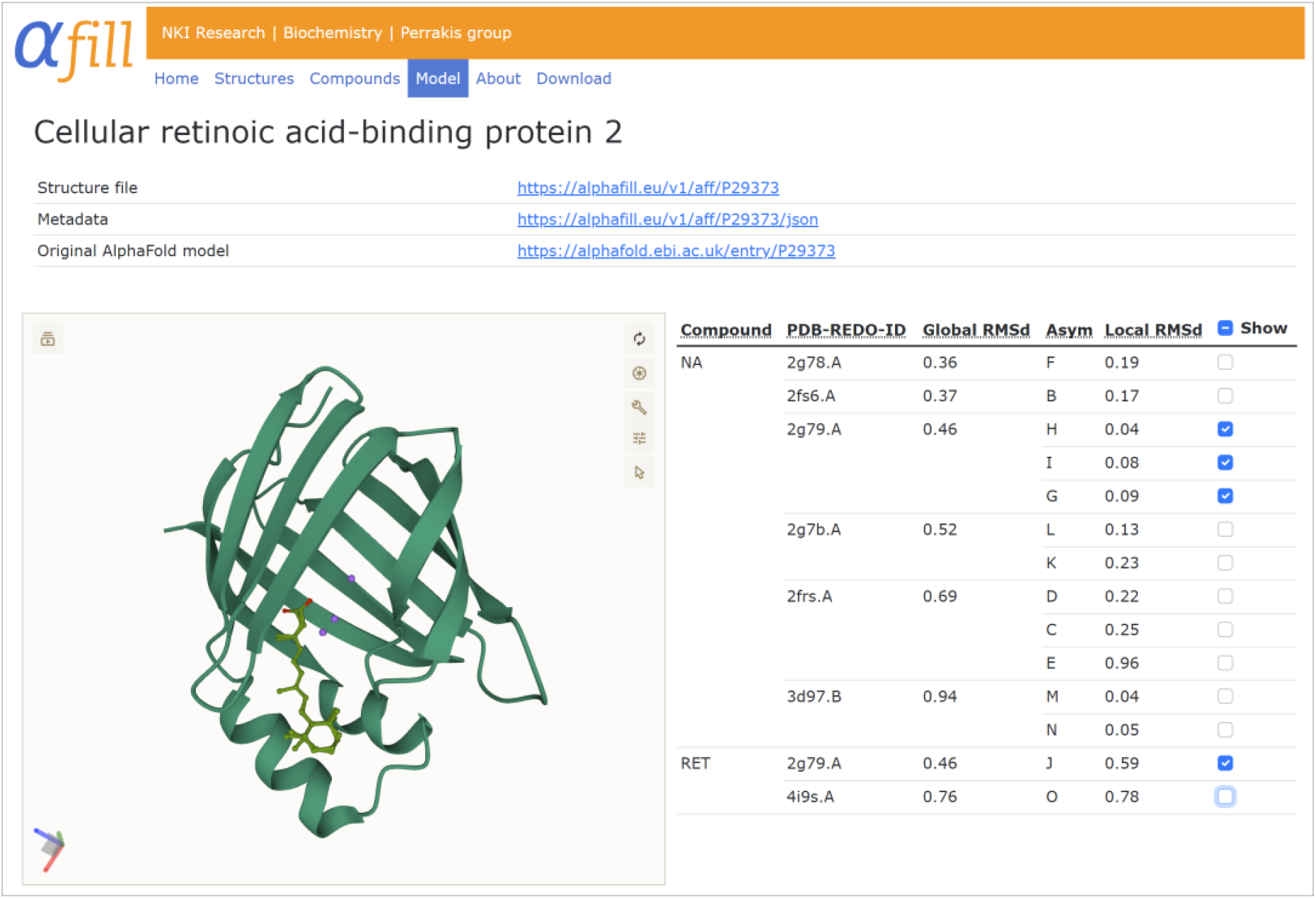
Example of the AlphaFill entry page with cellular retinoic acid-binding protein 2 (AF-P29373). The Mol* viewer on the left can be controlled by the list of transplanted compounds on the right. Clicking a compound brings up a zoom of the binding site. Compounds can be hidden or shown from view individually with the tick boxes. Here the retinal binding pose (with two alternate conformers) and three sodium ions inherited from PDB entry 2g79^14^ are shown. A distinct retinal binding pose from PDB entry 4i9s^15^ is hidden from view.

### Examples

The AlphaFill databank includes compounds transplanted based on identical experimentally determined protein structures, mapping to the same UniProt^16^ entry, and thus reproducing information such as already hyperlinked in the AlphaFold database to the PDBe-Knowledge Base^17^. In addition though, it transplants compounds from homologous structures that might have been determined in another species, and also to domains that have a high level of sequence and structure similarity to experimental structures. Thus, it offers additional functionality for the annotation of the models, that can help users make informed decisions about these structures. Here, we will discuss a few of such examples.

Human myoglobin is an ɑ-helical protein with heme B as cofactor, that can bind molecular oxygen and several other small molecules. The AlphaFold model (AF-P02144) is nearly identical to experimentally determined structures, but has a heme shaped cavity that can readily be supplemented with *AlphaFill* (Figure 2). The protein is well-studied with many heme analogues containing metals other than iron used in the crystallisation process, which are all mapped back to heme B (HEM, in PDB nomenclature). The heme analogues 6HE and 7HE, that exist as heme analogues in the CoFactor database, lack a carboxyl-tail and are therefore not mapped back to heme B, but are transferred as-is. It should be noted that the binding details of the AlphaFill model are suitable for qualitative analysis in terms of heme-protein interactions, but not for detailed quantitative analysis, such as measuring iron-histidine distances. Additional compounds that are transplanted to the AlphaFold model are molecular oxygen and carbon monoxide. The latter is fitted on two locations, one close to the iron atom in heme, and the other on the far side of the heme. Carbon monoxide at this unexpected location is inherited from PDB entry 1dwt^18^ in which it was modeled at 30% occupancy. This occupancy is retained in the AlphaFill model to allow users to take this into account when evaluating the model. The AlphaFill model of myoglobin also contains numerous metal ions. The cobalt and nickel ions that are shown, should be treated with care as they are inherited from engineered myoglobin dimers (PDB entries 7dgk^19^ and 7dgl^19^) that do not have a normal myoglobin fold. This is clearly reflected by the global alignment RMSD values being above 20Å, which is marked in the compound table by a question mark.

**Figure 2:**
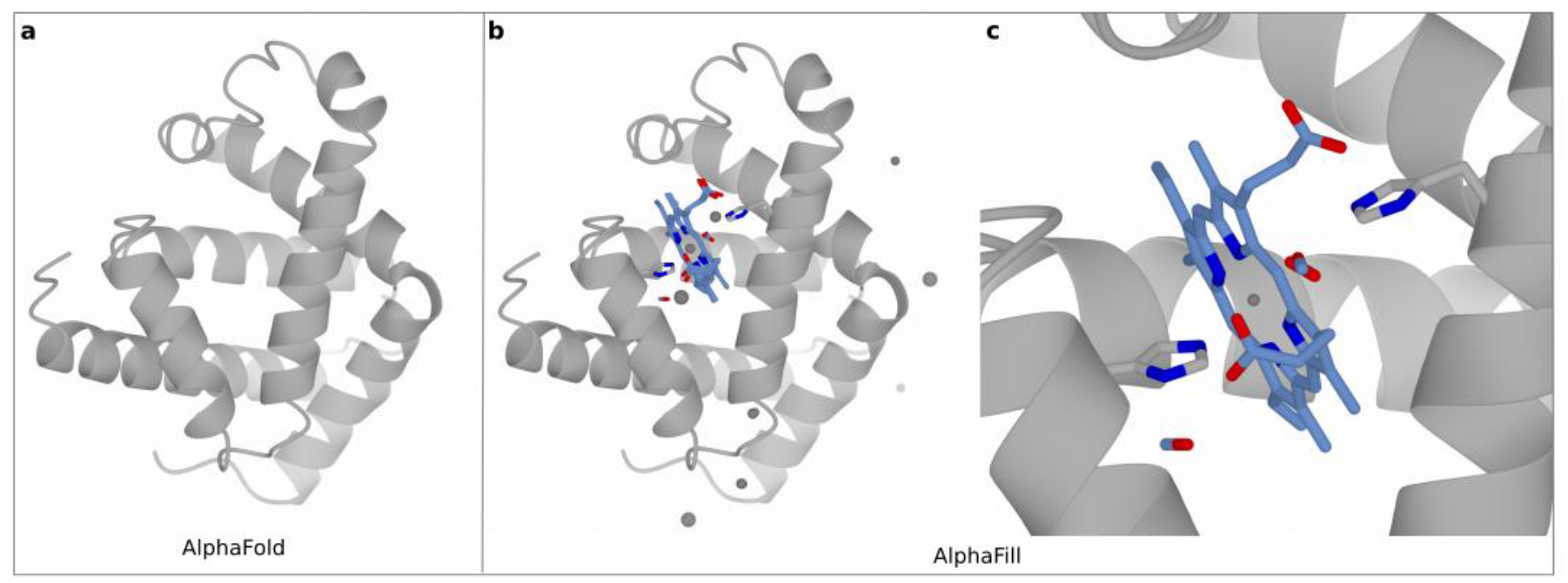
Human myoglobin structures in AlphaFold and AlphaFill. **a** A ribbon diagram of the AlphaFold model of human myoglobin. **b** The ribbon diagram of the AlphaFill model of myoglobin with all transplanted compounds; ligands are shown as blue cylinders coloured by atom type (heme) or spheres (ions); the histidine residues involved in coordinating the heme group are shown as grey cylinders coloured by atom type. **c** The heme group as in b; only the transplanted heme group (HEM) and the CO and O_2_ ligands are shown.

The most common transition-metal ion present in macromolecular structures is zinc (Table 1). Typically, it is involved in catalysis or in maintaining structural integrity^20^. The so-called “structural zinc ions” typically involve a tetrahedral binding site with a combination of four cysteine and/or histidine residues^21^. Such tetrahedrals are often distorted in the X-ray models available in the PDB, but the corresponding structures available through PDB-REDO contain improved binding sites^22^ and are a better reference for usage in *AlphaFill*.

The models provided in the AlphaFill databank in which a zinc ion was transplanted, can contain functional and/or structural zinc ions. One of the structures that contains both “types’’ of zinc ions is the structure of the STAM-binding protein, a zinc metalloprotease that cleaves lysine-63-linked polyubiquitin chains (AF-O95630)^23^. Zinc ions have been transplanted both at the catalytic site and at the zinc-finger motif (Figure 3a & b, respectively), originating from the PDB-REDO structure 3rzv^23^. The structural zinc ion is coordinated by three histidine residues and one cysteine. Although this tetrahedral zinc binding site looks proper, the atomic distances between the zinc atom and its ligands deviate substantially from previously established target values^22^. This limitation is a consequence of AlphaFold predicting the structure outside the context of key structural elements, in this case the zinc ions. By adding the zinc atom qualitative information is provided (the zinc atom should be in this binding site), but no quantitative information about the zinc binding site should be extracted from the AlphaFill model. Further refinement of the AlphaFill model with geometric restraints can be applied to make the binding site more normal, but this would not yield new information about zinc binding geometry.

**Figure 3:**
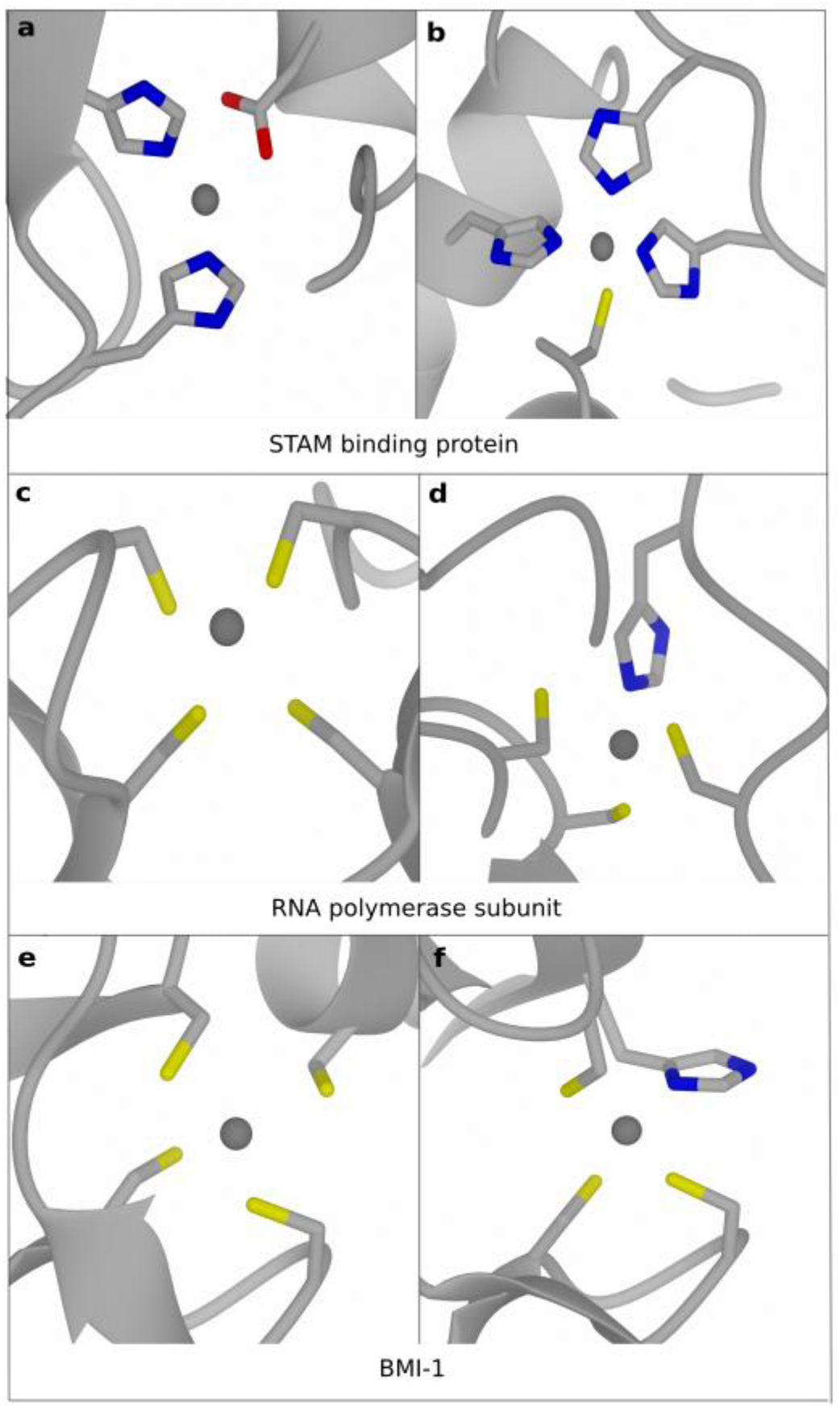
Examples of transplanted zinc ions (grey spheres) **a** & **b** A catalytic (a) and a structural (b) zinc ion in the STAM binding protein. **b** & **c** Structural zinc ions in the S. cerevisiae DNA-directed RNA polymerase II subunit RPB1 and **d** & **e** in the human BMI-1. Side chains coordinating the zinc ions are shown as grey cylinders coloured by atom type; all proteins are presented as a ribbon diagram and coloured in grey.

A similar situation is found for the two “transplanted” zinc ions in the DNA-directed RNA polymerase II subunit RPB1 of *Saccharomyces cerevisiae* (AF-P04050)^24^. One of the structural zinc ions in this structure model is coordinated by four cysteine residues (Figure 3c). The binding site is somewhat distorted in terms of coordination geometry with non-optimal coordination distances and cysteine side chain conformations, but the fact that this is a structural zinc site is very clear. This also holds for another zinc ion involved in the structural integrity of the same protein (Figure 3d). Both zinc ions were transplanted from PDB-REDO entry 1k83^24^. Similar tetrahedral zinc binding sites are found in the AlphaFold prediction of the human BMI-1 protein (AF-P35226), which contains two zinc binding sites that are involved in structural integrity^25^. These two zinc atoms were transferred by AlphaFill (Figure 3e & f), completing the structural overview of BMI-1 with respect to structural integrity.

In the ectonucleotide pyrophosphatase/phosphodiesterase (ENPP) family of proteins a bi-metalic zinc site is important for catalysis^26^. A structural alignment of the catalytic domain of the PDB-REDO models of ENPP1-7 (Figure 4a), shows that the zinc atoms and the residues that coordinate them occupy highly similar positions in all family members. The AlphaFold predictions of the same proteins (AF-P22413, AF-Q13822, AF-O14638, AF-Q9Y6X5, AF-Q9UJA9, AF-Q6UWR7, AF-Q6UWV6 for ENPP1-7, respectively) show slightly more divergence, especially for one of the histidine residues coordinating the zinc atom to the right (Figure 4b). *AlphaFill* picks up the similarity between the AlphaFold and the PDB-REDO models and transplants both zinc ions into the protein models of ENPPs (Figure 4c). As one histidine has different rotamers in the AlphaFold prediction, which clearly should be a single rotamer as can be concluded from the experimental structure model models, the bimetallic zinc site in the AlphaFill model(s) needs some refinement before usage.

**Figure 4:**
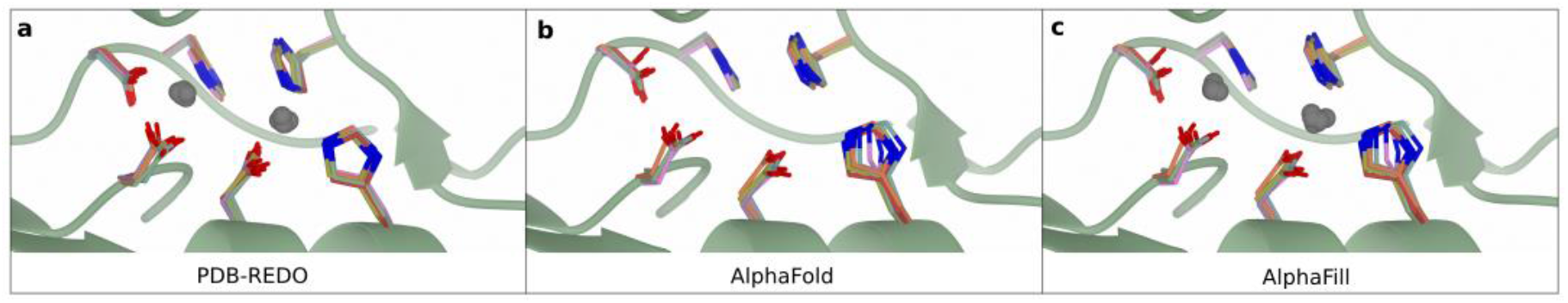
The bimetallic zinc binding site in ENPP1-7. **a** The bimetallic zinc binding site as found in PDB-REDO models (PDB identifiers for ENPP1-7: 6weu^27^, 5mhp^28^, 6c01^29^, 4lqy^30^, 5veo^31^, 5egh^32^, 5tcd^33^, respectively) **b** The same binding site as found in the human ENPP1-7 models from AlphaFold. **c** The bimetallic zinc binding site in the human ENPP1-7 models as available in AlphaFill, containing the two zinc ions. For clarity, only the backbone of ENPP1 is shown as a green ribbon diagram; side chains are coloured green, blue, red, pink, orange, purple, and gold for ENPP1-7 respectively.

The human heterodimer MSH2/MSH6 is involved in DNA mismatch repair. X-ray structures obtained for this dimer in complex with different DNA substrates and in the presence or absence of ADP show different conformations of both subunits for the apo structure versus the ADP bound state^34^. The experimental structure of the MSH2/MSH6 dimer in complex with a DNA containing a G-T mismatch, shows how the domains “clamp” the DNA strand, but also how the nucleotide bindings sites are formed with the participation of both monomers (Figure 5a, PDB-REDO entry 2o8b^34^). The two nucleotide binding sites have an ADP molecule and a magnesium ion, each (Figure 5b)^35^. Both subunits of this asymmetric ATPase are available through the AlphaFold database, albeit separately (MSH2: AF-P43246, MSH6: AF-P52701). Both the MSH2 and MSH6 models provided in the AlphaFill databank are “filled” with a magnesium ion and an ADP molecule (Figure 5c & d), based on the crystallographic models of the MSH2/MSH6 dimer that all contain bound ADP molecules (Figure 5a). Expectedly, both the predicted overall structure and the conformation of the nucleotide binding site are consistent with the ADP bound form that is experimentally available. The predicted MSH2/6 heterodimer can adopt multiple conformations depending on mismatch binding and (asymmetric) ATP hydrolysis, but that information is not captured in AlphaFold and cannot be enriched by *AlphaFill*. Interestingly though, the MSH2 subunit has an additional ADP molecule transferred by *AlphaFill*. This ADP comes from the X-ray model containing ADP and a G-dU mismatch (PDB-REDO entry 2o8d^34^), where it is bound to the MSH6 subunit. The backbone atoms of the 723-725 loop in that crystallographic structure of MSH2 are close enough to MSH6 that *AlphaFill* erroneously considers the ADP molecule as part of the MSH2 subunit, performing the local alignment and transplanting the molecule. This illustrates that the filled AlphaFold models should be analysed with care before usage. It is advised that the models from which the compounds are transferred are kept in mind during AlphaFill model analysis. This is why the parent structure models and chains are explicitly listed in the AlphaFill web interface and also provided in the metadata.

**Figure 5:**
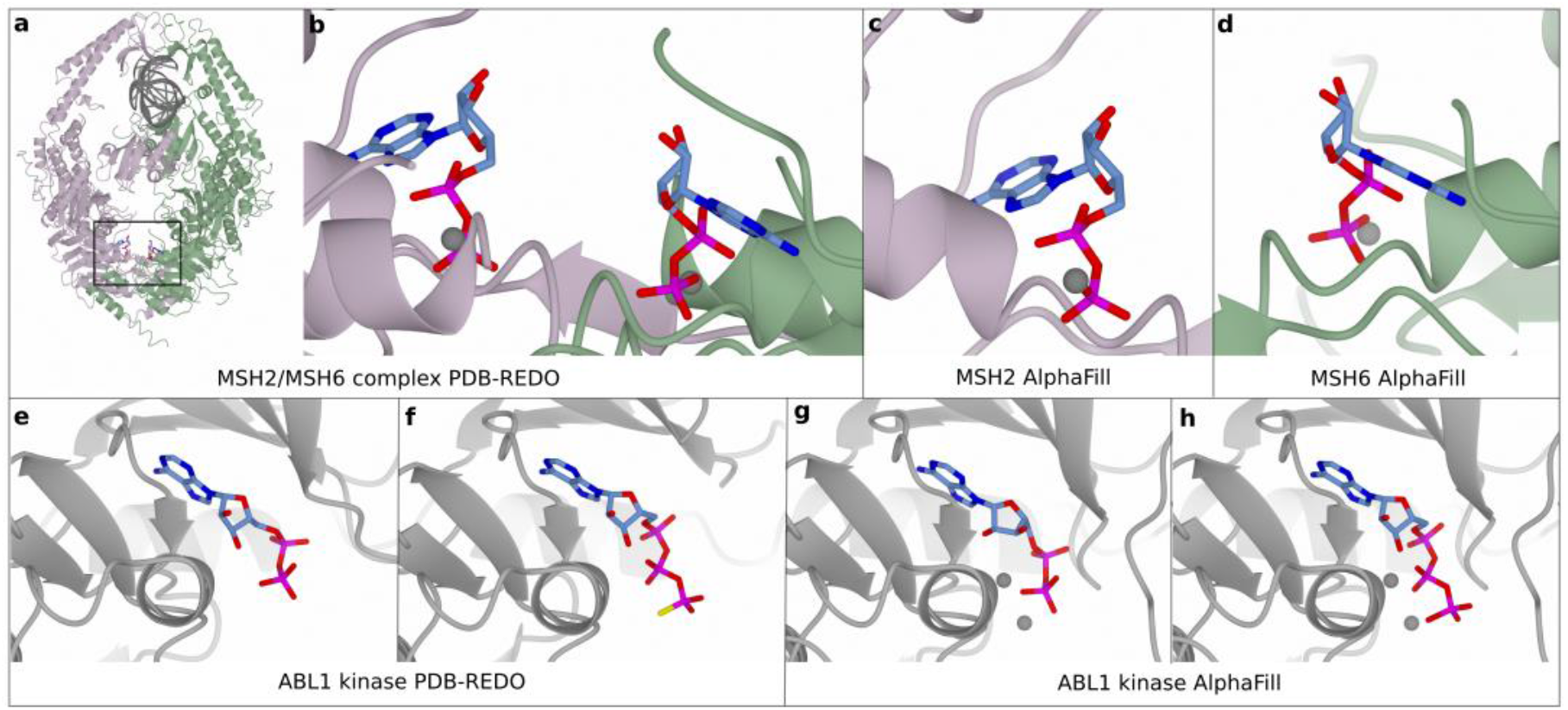
**a** PDB-REDO model of the human MSH2/MSH6 complex with two times ADP bound (PDB entry 2o8b), MSH6 in green ribbons, MSH2 in pink ribbons, DNA in grey ribbons, ions as grey spheres and ADP as cylinders coloured by atom type. **b** Zoom (black box of panel a) of the ADP binding site of the MSH2/MSH6 complex as in the PDB-REDO model. **c** MSH2 is filled with ADP and magnesium in AlphaFill model (AF-P43246). **d** MSH6 is filled with ADP and magnesium in AlphaFill model (AF-P52701). **e** ADP binding site of the human ABL1 kinase in PDB-REDO entry 2g2i which represents an active kinase state. **f** ABL1 kinase bound with AGS in PDB-REDO entry 2g2f which represents an “intermediate” kinase state. **g** AlphaFill model of the ABL1 kinase with ADP and magnesium ions shown (AF-P00519). The state of the kinase is not known a priori. **h** AlphaFill model of the ABL1 kinase with ATP (mapped from AGS) bound (AF-P00519).

Kinases are known to have multiple states, the extremes of which are the active conformation that offers an environment conducive to the phosphotransfer reaction, and the inactive state which does not fulfil the chemical constraints required for catalytic activity^36^. AlphaFold predicts only one conformation of a protein in which the state is not known *a priori*. *AlphaFill* transfers both ATP and ADP (or their analogues), to the AlphaFold model, provided that related experimental structures are available in the PDB-REDO databank. Importantly, *AlphaFill* will transfer ATP/ADP regardless of the state of the kinase, which is characterised by the conformation of specific residues. Nevertheless, the state of the kinase in the AlphaFold model can be partially evaluated based on the global and local alignment RMSD values. For the human tyrosine-protein kinase ABL1 (AF-P00519) the AlphaFold model is filled with an ADP molecule and an ATP molecule (Figure 5g & h), allowing in principle a hypothesis for the functional state represented in this model. The global RMSD for ADP is 2.54Å, and for ATP 1.36Å, while the local RMSD for ADP is 0.99Å, and for ATP 0.65Å, suggesting that the structure is more representative to the ATP-bound state. The interface informs the users that the ATP molecule was inherited from the “B” chain of the experimental structure 2g2f with bound AGS (ATP-γ-S) (Figure 5f), an ATP analogue whose binding promotes an “intermediate” state^37^. The ADP has been transplanted from PDB entry 2g2i^37^ (Figure 5e), which in this case represents an active state. Thus, the AlphaFill interface correctly indicates the differences, and allows a simple lookup of the PDB-models and the associated literature to draw relevant conclusions.

## Discussion

Cofactors, ligands, and ions help to understand both the function and structural integrity of proteins. They can also be helpful for designing downstream experiments, either computational or in the wet-lab. The AlphaFold database itself, recognises that need by offering links for each predicted model to experimental structures of the same protein, when they exist, by offering a link to PDBe-Knowledge Base. Here, we have described an algorithm used to create a public resource, that takes this further: we do not limit the mapping to identical sequences, but we extend it to homologues of this model.

Instead of “transplanting” all ligands that might occur in experimental structures for each AlphaFold model, we have chosen to limit our search to a list of common cofactors and common ions that are often found in protein structure and are likely to have a functional or structural role. This compound library was generated by human design and is limited by the number of occurrences in the PDB for ligands and by presence in the CoFactor database for the cofactors. As a consequence, *AlphaFill* will not fit compounds that are not annotated in this list, even though such compounds may be important in protein function or structure integrity. We have purposely chosen not to include certain compounds to this list. A notable example is glycosylation, for which the case has been already argued and exemplified for a handful of cases^38^. Glycosylation is a complicated issue that would need special attention, as would other post-translational modification - most notably phosphorylation which frequently induces conformational changes.

An important decision parameter in the AlphaFill algorithm is the minimum sequence identity threshold to allow transfer of information from an experimental structure to an AlphaFold model. We have chosen 70% identity as a limit in the BLAST search, based on our experience with homology restraints^7^ and homology-based annotation of experimental structures^9^. However, using more liberal thresholds could be argued, e.g. close to the 30% sequence identity limit that has been the threshold to obtain a reliable homology-based model for years.

Lowering the threshold too much though, might introduce to the transfer list paralogs from which ligands should not be transferred as the protein’s function is not (fully) conserved. To this end, we also note that currently, we use the global and the local RMSD as metadata that are presented to the user as indicators of the reliability of ligand modeling. It can also be argued that these can also be useful as thresholds for not “transplanting” a ligands, especially if lower sequence identity thresholds are utilised for ligands transfer in the near future. Additionally, defaults regarding the sequence alignment length (85 amino acids), the local structure alignment surrounding the “transferable” compound (6.0Å), and the distance between two “transferrable” compounds (3.5Å) were set. Investigation of these default parameters could also be examined further, as they also affect the outcomes.

The AlphaFill structure models are not meant to be accurate or precise or complete representations of the full repertoire of ligands for a certain protein structure. They are meant more as a tool for the non-expert. Most structural biology or structural bioinformatics experts, would find it trivial to select, superpose, and “transplant” a functional or structural ion (e.g. a zinc, as in our Figure 2 examples) and take that information for molecular dynamics simulations, mutagenesis studies, or discussing the structure of a model in light of new biochemical or biophysical insights. However, that would be less obvious for the non-experts. Here, we aim to provide a resource for all life scientists, to enrich the transformative impact that AlphaFold database models are already having on biomolecular research. We find that the AlphaFill databank, can offer easy access to “liganded” models (even for experts that would not need to manually perform the computation procedures that the *AlphaFill* algorithms have now automated), but also - very importantly - offer clear information on the provenance of the information we provide. To maximise the impact of the AlphaFill resource, it should ideally be linked directly to the AlphaFold database.

It is also good to keep in mind, that the AlphaFill models are not meant or suitable for precise quantification of interactions between the transferred ligand(s) and the protein (e.g. hydrogen bonds, π-π or cation-π interactions, van der Waals interactions, hydrophobic interactions, halogen bonds). These require coordinate precision that is not provided by either the AlphaFold or the AlphaFill models at the current stage, and the models should only be interpreted in a qualitative manner. It should be noted that in some cases ligand interactions come from parts of the protein that are not modeled with high confidence by AlphaFold. This confidence is presented as the per-residue pLDDT value^1^ which is stored in the model files in the *_ma_qa_metric_local.metric_value* records. The pLDDT value can also be used as a reference to assess the reliability of perceived protein-ligand interactions. It should also be noted that future work should aim to increase the accuracy of the AlphaFill - and thus the AlphaFold models - by optimising the local environment of important and/or accurately predicted ligands.

The *AlphaFill* software is made to be flexible by design, so that the settings used in this study can easily be altered. Similarly, the list of compounds can readily be updated based on user requirements. As always, the development of scientific resources depends on constructive user feedback and as such this is also appreciated for AlphaFill. Future incarnations of the basic idea should be envisaged in the context of different - or perhaps dynamic - thresholds for decision making. To that end, AlphaFill by definition depends on homology as the first and major criterion for transferring ligands. However, it is well established that certain structural motifs can occur outside the context of sequence similarity. AlphaFill should be complemented by structure-based transfer algorithms, likely based on deep learning concepts similar to those used for AlphaFold structure prediction revolution, that could further enrich our understanding of AlphaFold structures enriching them with ligands that cannot be currently predicted based on sequence information alone, or not even based on structural similarity searches (e.g. DALI^39^ and PDBeFold^40^).

## Methods

### Input data

All available AlphaFold models^1^ were downloaded from the AlphaFold Protein Structure Database’s FTP archive. A local copy of the PDB-REDO databank^7^ was used to provide ligands for transfer.

To find all relevant PDB-REDO entries for a specific AlphaFold model through sequence-based retrieval with BLAST^8^ a PDB-REDO-specific sequence database (as of November 1st 2021) was used. This database is created automatically as part of the weekly LAHMA^9^ and PDB-REDO databank updates.

The selection of biological relevant molecules to be added to the AlphaFold models was performed based on the number of occurrences in the PDB. All ions and small molecule ligands occurring over 1600 times were added to the initial AlphaFill compound list, as well as all cofactors and cofactor analogues present in the organic CoFactor database^12^. To map cofactors analogues and cofactor adducts to their canonical cofactors (e.g. non-hydrolysable ATP-analogue ACP to ATP), equivalent atoms have to be renamed or removed. The required changes were found by visual inspection of the compounds via the Ligand-Expo website^41^ and the PDB web sites and stored in a cif-formatted data file that can easily be extended. Cofactor analogues that have atoms missing with respect to their parent are kept as-is. From the AlphaFill ligand list, common crystallisation agents (e.g. poly-ethyleneglycol and chloride), compound used for phasing (e.g. cadmium ions) and, carbohydrates and biologically non-relevant ligands were removed based on manual curation. The final compound list contains 400 compounds.

### The *AlphaFill* software

A new program, *AlphaFill*, was created for the purpose of this study. The program reads an AlphaFold model together with the AlphaFill compound list and the PDB-REDO-specific sequence database and returns an AlphaFill model consisting of the AlphaFold model coordinates plus all transferred compounds. See the main text for the compound transfer procedure. The *AlphaFill* program is based on the *libzeep*^42^, *libcif++* (a general purpose C++ library for dealing with mmCIF data structures), *libpdb-redo* (a core library for PDB-REDO software) and *clipper*^43^ libraries and contains its own BLAST implementation. The source codes of *AlphaFill*, *libcif++*, and *libpdb-redo* are available from https://github.com/PDB-REDO.

### Creation of the AlphaFill databank

The AlphaFill databank was created by running *AlphaFill* over all AlphaFold models. Consistent with other structure-derived databanks, this was done using the *make* system with the AlphaFold coordinate files as sources and the AlphaFill coordinate files as targets^44^. The calculation was *embarrassingly parallel* and took 10 days and 14.5 hours using 48 CPU threads on Intel Xeon E5-2697 CPUs @ 2.30GHz.

### The AlphaFill web interface

The web site is created as a web application using the libzeep library which offers an HTTP(S) server, HTML templating and many other components for web server construction in C++. Handling of mmCIF files is done using libcifpp. The model is presented on the page using Mol*^13^ as an interactive web component.

### Model analysis

The AlphaFill models were analysed visually using Coot^45^, the AlphaFill website and CCP4mg^46^. Molecular graphics figures were made with CCP4mg^46^.

## Acknowledgements

The authors thank the Research High Performance Computing facility of the Netherlands Cancer Institute for providing and maintaining computation resources and Dr. Stuart McNicholas for support with CCP4mg. This work has been supported by iNEXT-Discovery, project number 871037, funded by the Horizon 2020 program of the European Commission.

## Author Contributions

MLH developed the AlphaFill software and web interface. IdV analysed chemical compounds for integration, worked on data FAIRification, and prepared the example cases and related figures. AP and RPJ conceived and supervised the project. All authors contributed to the manuscript, the experimental and algorithmic design, and the analysis of the results.

