## Supplemental Table S1 for "AlphaFill: enriching the AlphaFold models with ligands and co-factors"

**Supplemental Table 1: List of Compounds.** This table presents the compounds with their analogue and the number of AlphaFold models to which the compound was transplanted (# of structures), plus the number on how often the compound was transplanted in total (# of transplants) by *AlphaFill*.

| Compound | Analogue | # of structures | # of transplants |
|---|---|---|---|
| MG | MG | 8069 | 19754 |
| NA | NA | 5650 | 15262 |
| ZN | ZN | 5261 | 13051 |
| CA | CA | 5078 | 17312 |
| K | K | 2267 | 3910 |
| ADP | ADP | 2198 | 2928 |
| MN | MN | 1814 | 3973 |
| GDP | GDP | 1655 | 2468 |
| ANP | ATP | 1358 | 1741 |
| NI | NI | 1249 | 2418 |
| ATP | ATP | 1221 | 1517 |
| NO3 | NO3 | 776 | 2873 |
| AMP | AMP | 723 | 926 |
| CO | CO | 695 | 1290 |
| GTP | GTP | 686 | 921 |
| NAD | NAD | 685 | 978 |
| GNP | GTP | 633 | 721 |
| HEM | HEM | 498 | 658 |
| FE | FE | 472 | 746 |
| FAD | FAD | 470 | 628 |
| NAP | NAP | 455 | 618 |
| PLP | PLP | 369 | 602 |
| SAH | SAH | 355 | 383 |
| GSH | GSH | 324 | 450 |
| NDP | NDP | 317 | 445 |
| SAM | SAM | 298 | 407 |
| AGS | ATP | 287 | 370 |
| ACP | ATP | 284 | 411 |
| COA | COA | 254 | 368 |
| FE2 | FE2 | 248 | 371 |
| UDP | UDP | 243 | 267 |
| CU | CU | 217 | 439 |
| NAI | NAI | 204 | 298 |
| FMN | FMN | 197 | 266 |

| Compound | Analogue | # of structures | # of transplants |
|---|---|---|---|
| CO3 | CO3 | 195 | 323 |
| POP | POP | 192 | 275 |
| ACO | COA | 170 | 203 |
| CLA | CLA | 163 | 4058 |
| CLR | CLR | 157 | 380 |
| OGA | AKG | 144 | 150 |
| OXY | OXY | 143 | 214 |
| SF4 | SF4 | 128 | 172 |
| FES | FES | 121 | 147 |
| TAD | TAD | 112 | 112 |
| BCR | BCR | 93 | 442 |
| SFG | SFG | 91 | 94 |
| DTP | DTP | 85 | 146 |
| HEC | HEC | 76 | 209 |
| XAT | XAT | 75 | 114 |
| CMO | CMO | 69 | 81 |
| TPP | TPP | 64 | 92 |
| SOP | COA | 60 | 60 |
| PMP | PMP | 55 | 93 |
| CHD | CHD | 52 | 155 |
| THG | THG | 36 | 54 |
| MDO | MDO | 35 | 35 |
| PLR | PLR | 34 | 49 |
| FDA | FDA | 32 | 35 |
| C2F | C2F | 31 | 47 |
| HBI | HBI | 30 | 35 |
| ASC | ASC | 30 | 33 |
| B12 | B12 | 26 | 31 |
| RET | RET | 26 | 37 |
| COM | COM | 25 | 41 |
| CAA | COA | 24 | 31 |
| GDS | GDS | 24 | 42 |
| F3S | F3S | 24 | 29 |
| PL9 | PL9 | 24 | 67 |
| AT5 | AT5 | 19 | 19 |
| MTE | MTE | 19 | 19 |
| UQ1 | UQ1 | 17 | 25 |
| SRM | SRM | 17 | 17 |
| MH0 | MH0 | 16 | 16 |
| FNR | FNR | 16 | 17 |

| Compound | Analogue | # of structures | # of transplants |
|---|---|---|---|
| H4B | H4B | 16 | 33 |
| GPS | GSH | 16 | 19 |
| PHO | PHO | 15 | 31 |
| FFO | FFO | 15 | 26 |
| ABY | ABY | 14 | 28 |
| GTX | GSH | 14 | 26 |
| SND | NAD | 14 | 14 |
| TZD | TZD | 14 | 14 |
| PQN | PQN | 14 | 14 |
| HMG | COA | 13 | 13 |
| DCA | DCA | 13 | 13 |
| BOB | GSH | 13 | 14 |
| TAP | NAD | 13 | 13 |
| CMC | COA | 12 | 12 |
| MYA | COA | 12 | 24 |
| 01K | COA | 11 | 11 |
| FAB | FAB | 11 | 11 |
| GBX | GSH | 11 | 20 |
| RBF | RBF | 11 | 13 |
| TXE | NAD | 11 | 12 |
| FRE | COA | 10 | 10 |
| HXC | COA | 10 | 14 |
| MLC | COA | 10 | 11 |
| BTN | BTN | 9 | 12 |
| LZ6 | GSH | 9 | 16 |
| BTI | BTI | 9 | 15 |
| DTB | DTB | 9 | 10 |
| SAE | SAE | 9 | 9 |
| THV | THV | 8 | 8 |
| TDM | TDM | 8 | 8 |
| LPA | LPA | 8 | 9 |
| 6V0 | 6V0 | 7 | 7 |
| 18W | 18W | 7 | 7 |
| COF | COA | 7 | 7 |
| T6F | T6F | 7 | 9 |
| HAX | COA | 7 | 7 |
| COD | COD | 7 | 9 |
| GSM | GSH | 7 | 7 |
| NAJ | NAJ | 7 | 12 |
| 4IK | SAM | 6 | 6 |

| Compound | Analogue | # of structures | # of transplants |
|---|---|---|---|
| NHW | COA | 6 | 6 |
| NHQ | COA | 6 | 12 |
| F43 | F43 | 6 | 8 |
| TGG | GSH | 6 | 6 |
| GTD | GSH | 6 | 6 |
| COO | COA | 6 | 12 |
| HEA | HEA | 6 | 10 |
| 3GC | 3GC | 6 | 11 |
| PP9 | PP9 | 6 | 14 |
| SMM | SAM | 5 | 5 |
| FA8 | FA8 | 5 | 5 |
| DPM | DPM | 5 | 5 |
| HEB | HEB | 5 | 9 |
| BCO | COA | 5 | 5 |
| CO8 | COA | 5 | 6 |
| GPR | GSH | 5 | 9 |
| UQ5 | UQ5 | 5 | 11 |
| UQ6 | UQ6 | 5 | 5 |
| MTQ | MTQ | 5 | 5 |
| H2B | H2B | 5 | 5 |
| WCA | COA | 5 | 5 |
| TP7 | TP7 | 5 | 7 |
| TT8 | TT8 | 4 | 4 |
| FON | FON | 4 | 6 |
| GBP | GSH | 4 | 4 |
| GS8 | GSH | 4 | 5 |
| GSB | GSH | 4 | 10 |
| GSF | GSH | 4 | 6 |
| GTS | GSH | 4 | 7 |
| CO6 | COA | 4 | 4 |
| CNC | B12 | 4 | 4 |
| CAO | COA | 4 | 8 |
| AHE | GSH | 4 | 8 |
| 8ID | NAD | 4 | 8 |
| NBP | NAD | 4 | 4 |
| 6NR | 6NR | 4 | 4 |
| 5AU | 5AU | 4 | 8 |
| PNS | PNS | 4 | 10 |
| 1U0 | TPP | 4 | 4 |
| THF | THF | 4 | 8 |

| Compound | Analogue | # of structures | # of transplants |
|---|---|---|---|
| THH | THH | 4 | 5 |
| THW | THW | 4 | 4 |
| THY | THY | 4 | 4 |
| 0XU | SAM | 4 | 4 |
| WWF | WWF | 4 | 4 |
| 0WD | NAD | 4 | 4 |
| FDE | FDE | 4 | 4 |
| 6HE | 6HE | 3 | 3 |
| MGD | MGD | 3 | 4 |
| 7HE | 7HE | 3 | 3 |
| XAX | XAX | 3 | 3 |
| 4YP | 4YP | 3 | 3 |
| 76H | SAM | 3 | 3 |
| SCA | COA | 3 | 9 |
| ZNH | HEM | 3 | 3 |
| 1VU | COA | 3 | 3 |
| HSC | COA | 3 | 3 |
| GNB | GSH | 3 | 5 |
| 37H | SAM | 3 | 3 |
| T5X | T5X | 3 | 5 |
| ZOZ | COA | 3 | 3 |
| TXP | NAD | 3 | 3 |
| ZBF | GSH | 3 | 6 |
| MQ7 | MQ7 | 3 | 5 |
| BHS | BHS | 3 | 6 |
| GSO | GSH | 3 | 6 |
| 1R4 | GSH | 3 | 3 |
| BIO | BIO | 2 | 4 |
| COZ | COA | 2 | 2 |
| TS5 | TS5 | 2 | 2 |
| GTB | GSH | 2 | 3 |
| HAG | GSH | 2 | 2 |
| CUA | CUA | 2 | 2 |
| TD6 | TD6 | 2 | 4 |
| MCN | MCN | 2 | 2 |
| HTL | HTL | 2 | 4 |
| COH | HEM | 2 | 2 |
| TXZ | TXZ | 1 | 2 |
| 8FL | 8FL | 1 | 1 |
| 0Y0 | 0Y0 | 1 | 1 |

| Compound | Analogue | # of structures | # of transplants |
|---|---|---|---|
| NDE | NAD | 1 | 1 |
| 8EO | 8EO | 1 | 1 |
| BYC | COA | 1 | 1 |
| MDE | COA | 1 | 1 |
| 8EL | 8EL | 1 | 1 |
| 8EF | 8EF | 1 | 1 |
| ESG | GSH | 1 | 1 |
| UQ2 | UQ2 | 1 | 1 |
| DCC | COA | 1 | 2 |
| 4LU | FMN | 1 | 1 |
| PXP | PXP | 1 | 1 |
| R1T | R1T | 1 | 1 |
| COB | B12 | 1 | 1 |
| S0N | COA | 1 | 2 |
| SA8 | SAM | 1 | 1 |
| HDE | HEC | 1 | 2 |
| 3CD | NAD | 1 | 2 |
| 3AA | 3AA | 1 | 1 |
| 2NE | COA | 1 | 3 |
| HDD | HDD | 1 | 2 |
| 1XE | COA | 1 | 1 |
| 1JO | GSH | 1 | 1 |
| TD7 | TD7 | 1 | 2 |
| TD8 | TD8 | 1 | 2 |
| TD9 | TD9 | 1 | 2 |
| TDK | TDK | 1 | 2 |
| TDW | TDW | 1 | 2 |
| 0ET | COA | 1 | 4 |
| 1HA | COA | 1 | 1 |
| ODP | NAD | 1 | 1 |
| FAO | FAD | 1 | 2 |
| 1DG | NAD | 1 | 1 |
| TPW | TPW | 1 | 1 |
| TPZ | TPZ | 1 | 2 |
| FMI | HEC | 1 | 1 |
| MNR | HEM | 1 | 1 |
